# Epicardial transplantation of autologous atrial appendage micrografts–evaluation of safety and feasibility in pigs after coronary artery occlusion

**DOI:** 10.1101/2021.10.18.464894

**Authors:** Annu Nummi, Tommi Pätilä, Severi Mulari, Milla Lampinen, Tuomo Nieminen, Mikko I. Mäyränpää, Antti Vento, Ari Harjula, Esko Kankuri, the AADC consortium

## Abstract

Several approaches devised for clinical utilization of cell-based therapies for heart failure often suffer from complex and lengthy preparation stages. Epicardial delivery of autologous atrial appendage micrografts (AAMs) with a clinically used extracellular matrix (ECM) patch provides a straightforward therapy alternative. We evaluated the operative feasibility and the effect of micrografts on the patch-induced epicardial foreign body inflammatory response in a porcine model of myocardial infarction. Right atrial appendages were harvested and mechanically processed into AAMs. The left anterior descending coronary artery was ligated to generate acute infarction. Patches of ECM matrix with or without AAMs were transplanted epicardially onto the infarcted area. Four pigs received the ECM and four received the AAMs patch. Cardiac function was studied by echocardiography both preoperatively and at three weeks follow-up. The primary outcome measures were safety and feasibility of the therapy administration and the secondary outcome was the inflammatory response to ECM. Neither AAMs nor ECM patch-related complications were detected during the follow-up time. AAMs patch preparation was feasible according to time and safety. Inflammation was greatly reduced in AAMs as compared to ECM patches as measured by the amount of infiltrated inflammatory cells and area of inflammation. Immunohistochemistry demonstrated an increased CD3+ cell density in the AAMs patch infiltrate. Epicardial AAMs transplantation demonstrated safety and clinical feasibility. The use of micrografts significantly inhibited ECM-induced foreign body inflammatory reactivity. Transplantation of AAMs shows good clinical applicability as adjuvant therapy to cardiac surgery and can suppress acute inflammatory reactivity.

## Introduction

Myocardial infarction (MI) is the major cause of death worldwide [1]. It bears a poor prognosis particularly when accompanied with ischemic heart failure. Revascularization and medical therapy are critical for severe ischemic heart failure but the recovery of the irreversibly damaged infarction area is poor or nonexistent. Cell therapy has been proposed as an additional strategy to the current treatment modalities for heart failure: one with potential to restore or even regenerate structure and function of the infarcted area. Several studies have suggested benefit from cell therapies, but therapy preparation suffers from complex and lengthy protocols, and the treatment requires additional interventions. Cell therapies exert their benefit largely through secretion of soluble paracrine factors. [2, 3] These protective factors activate pathways in the target tissue that result in the repair and remodeling of infarcted myocardium. [4, 5] As adjuvant to cardiac surgery, cell therapy holds potential to reduce mortality and re-hospitalization caused by heart failure during long term follow-up, improve global left ventricular ejection fraction (LVEF), New York Heart Association (NYHA) - functional class and quality of life as well as to lower poor prognosis-associated high NT-proBNP levels. [6] In our earlier studies the use of injected bone marrow mononuclear cells during coronary artery bypass (CABG) surgery showed significant reduction of myocardial infarction scar [7, 8], which is a major prognostic factor for survival in ischemic heart failure [9, 10].

Atrial appendage offers a good tissue-matched reservoir for various cell types, including progenitor cells, contributing to paracrine healing. [11–13] Moreover, it is thought that stem or progenitor cells of cardiac origin are more likely to differentiate into cardiac lineages [14] and may thus contribute to the atrial appendages’ effect on the failing myocardium. Autologous cells can minimize rejection and thus ensure better cell engraftment. The extracellular matrix and the microtissue architecture of the micrografts can support cellular adherence and survival of transplanted cells. [15] Moreover, the mixture of different myocardial cell types can enable better interplay and therapeutic effect via enhanced survival and improved paracrine signaling. [16, 17]

The effect of any cell therapy is critically dependent on the delivery method. [18] Animal models have proven the epicardial delivery route to be beneficial in securing generous cell engraftment when compared to various types of cell delivery routes by intramyocardial injection or coronary infusion. [19, 20] In principle, the technique of delivering progenitor cells epicardially causes minimal harm to the functional myocardium, less arrhythmogenicity [21] and ensures sufficient amount of cells to remain at the transplant area [18].

We have encouraging results on the transplantation of epicardial atrial appendage micrografts (AAMs) in ischemic heart failure in rodents [22] and from a clinical safety and feasibility trial in patients during CABG surgery [23]. This study was established to test the critical technical questions regarding the procedure and ensure the general safety. Specific focus was on the technical details of preparing the transplant (including sterility and time schedule/delay for preparing the transplant during open heart surgery). Additionally, we evaluated the effect of the micrografts on the foreign body reaction and inflammation caused by one of the clinically used pericardial patch extracellular matrix (ECM) sheets in this pig model.

## Materials and methods

All procedures on laboratory animals and animal care were approved by the Division of Health and Social Services, Legality and Licensing of the Regional State Administrative Agency for Southern Finland (ESAVI/1482/04.10.07/2015). The study protocol was approved by the Surgical Ethics Committee of the Hospital District of Helsinki and Uusimaa (number 180/13/03/02/13).

The study included eight pigs. Four animals received an ECM patch and four animals an ECM patch with AAMs. The procedure and follow-up were done similarly to all animals.

A standard anterior sternotomy was performed under anesthesia. First echocardiography (echo) was done prior to any changes in cardiac function. Then the right atrial appendage (RAA) was ligated using a purse string suture. RAA was removed from all animals in both groups. For the animals in the AAMs patch group, the harvested RAA was processed mechanically on-site in the operating room using a tissue homogenizer (Rigenera-system, HBW s.r.l., Turin, Italy) [24]. This system utilizes a sterile, single-use tissue homogenizer surface to generate the micrografts and yields approximately 5–10 millions of viable cells per gram of RAA tissue.

Myocardial infarction was introduced by ligating the left anterior descending coronary artery (LAD) with a non-absorbable suture (**Fig 1**). Optimal location for ligation was chosen in the B-part of the LAD so that the biggest diagonal artery remained untouched and open to secure adequate blood flow to the anterior wall of the left ventricle (LV). This was to make sure that the infarction was large enough to cause observable heart failure but the risk for fatal ventricular arrhythmias was controlled. The acute infarction was verified with reduced motion of the inferior wall, accelerated heart rhythm and the local changes in myocardial color.

**Figure 1.**
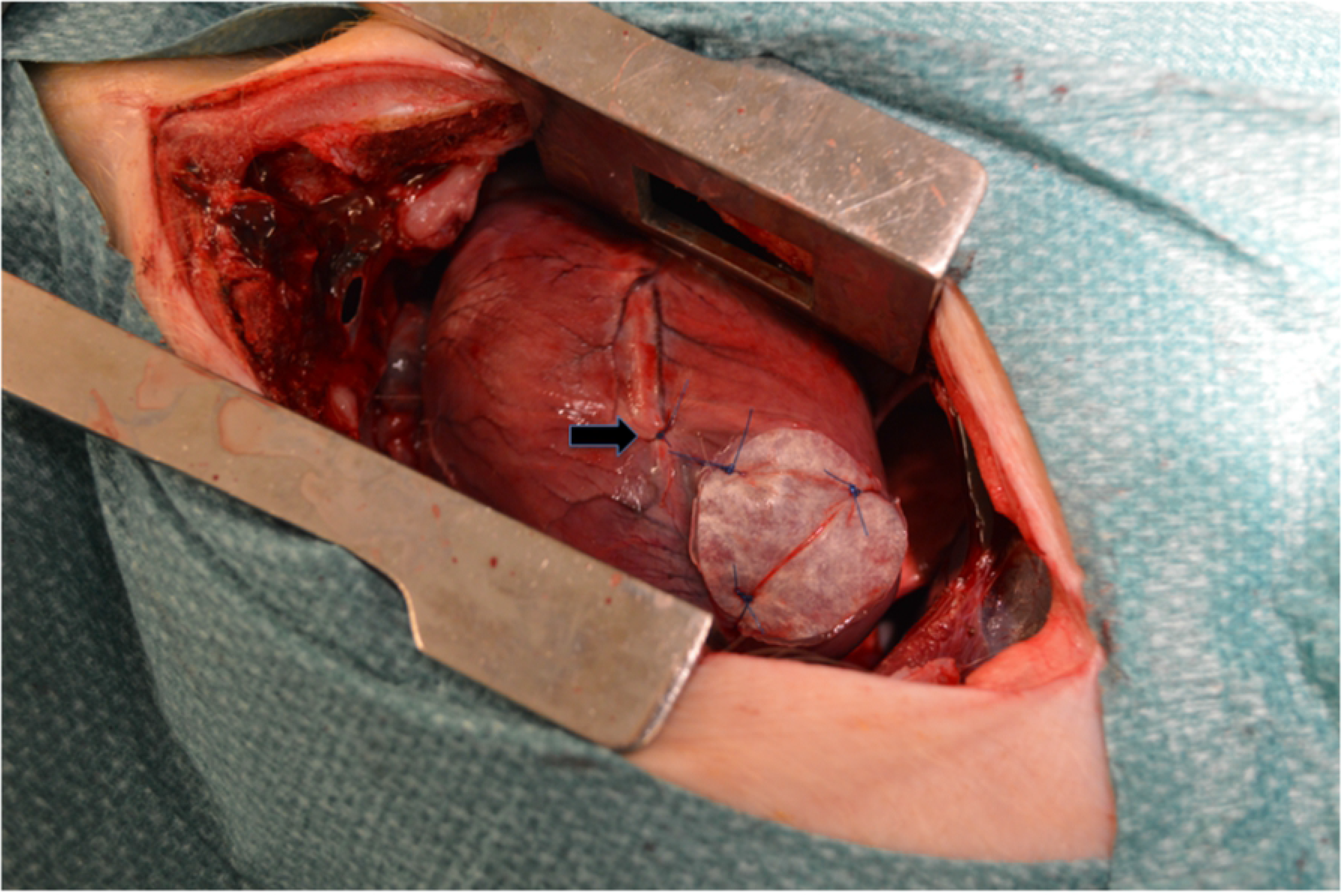
Completed patch at the area of infarct. Completed autologous atrial appendage micrograft (AAM) patch secured to myocardium with three sutures and ligation of left descending coronary artery causing heart infarct and failure (marked with black arrow).

The isolated AAMs were applied in standard cardioplegia suspension on the myocardium using the epicardial transplantation technique. We used an extracellular matrix sheet (Cormatrix^**®**^ ECMTM Technology, Cormatrix Cardiovascular Inc., Atlanta, GA, USA) where the suspension of AAMs were applied using fibrin sealant (Tisseel^TM^, Baxter Healthcare Corp. Westlake Village, CA, USA). Approximately two hours after producing the infarct and careful follow up of arrhythmias, the Cormatrix^®^ with the cell suspension was secured to the myocardial surface by three to four simple knots using non-absorbable suture. The detailed preparation of the cell sheet has been explained in our previous publication. [25]

Echo was performed by a single cardiologist. This was done under anesthesia and on open heart to achieve adequate vision. Echo was performed twice each animal; in the beginning of the first operation, prior to the atrial appendectomy and infarction, and at three weeks follow up when the animals were first put under anesthesia and later euthanized. Echo was done to measure the dimensions of the anterior wall of LV, to determine the changes in LVEF and to observe any changes in the cardiac function.

After the operation animals received painkillers (buprenorphine 0,3mg x2-3 and carprofen 50 mg x1-2) for three days. At the three weeks’ time point each animal was euthanized and samples from the infarction area with the patch was taken for histological analysis.

Hematoxylin-eosin (HE) staining and immunohistochemistry for CD45, CD3, and CD20 were performed and the plates were scanned by Genome Biology Unit (Biomedicum, University of Helsinki, Helsinki, Finland). Digital images were analyzed using Pannoramic Viewer (3DHISTECH Ltd, Budapest, Hungary) and MIPAR^™^ image analysis software (Worthington, OH, USA) was used for cell counting. Distance from Cormatrix^®^ inner surface to epicardium was measured at ten points from the distant corners of the patch in both groups. At the same distances, squared image samples of 1 mm^2^ were taken and MIPAR^™^ was used to calculate number of nuclei and their relative distances in both groups.

The primary antibodies were rabbit anti-CD3 IgG (SP7 monoclonal, Spring Bioscience M3072, Abcam, Cambridge, UK), rabbit anti-CD20 IgG (polyclonal, Thermo Fisher Scientific Labvision RB9013P, Waltham, MA, USA), rabbit anti-CD45 IgG (clone #145 monoclonal recombinant, Sino Biological 100342-R145, Beijing, China). All antibodies were diluted using the antibody diluent (BiositeHisto, Nordic Biosite, Tampere, Finland, cat. no BCB-QUG2XK). Secondary antibody was an HRP-polymer anti-rabbit antibody (BiositeHisto Nordic Biosite cat. no KDB-Z47C3W). Immunoreactivity of antibodies was controlled in sections of porcine kidney, spleen and liver using human tonsilla as positive control.

Immunohistochemistry was performed using the LabVision Autostainer 480 (Thermo Fisher Scientific) and heat-induced epitope retrieval for 20min at 98°C, followed by wash (TBS-TWEEN pH8,4), incubation with primary antibody (30min, RT), wash, detection polymer incubation (30min, RT), endogenous peroxidase blocking (10 min, H2O2), washx2, DAB (High Contrast DAB, BiositeHisto Nordic Biosite cat. no BCB-R7IKBJ, 10min, RT), wash, 0,5% CuSO4-enhancement (10min), wash, 1:10 Mayer’s hemalum solution (Merck KGaA, Darmstadt, Germany, 2min), bluing with running tap water (7min), and finally followed by dehydration-clearing. Mounting of coverslips was carried out with xylene-based mounting medium.

Slides were digitalized as WSI in Mirax format with 3DHistech Pannoramic MIDI scanner (Budapest, Hungary) at a pixel size of 0.23μmx0.23μm. The scanner utilizes a brightfield microscope setup with an HV_F22CL camera (Hitachi Kokusai Electric America Ltd, Southwick, MA, USA) equipped with a plan-apo 20x objective. For immunostaining analysis, serial non overlapping images covering the Cormatrix area from every sample were captured with the Panoramic Viewer software (3DHISTEC Ltd). Image area was then standardized as mm^2^. The analysis of the captured images was carried out using the FiJI ImageJ software. [26] The image analysis macros are available upon request from the corresponding author. Briefly, for each image, after background subtraction, the color deconvolution algorithm to hematoxylin (H) and diaminobenzidine (DAB) channels was utilized. The DAB channel image was thresholded to the stain using the automated default method based on the IsoData algorithm, and the stain intensity was measured. For nuclear counting, hematoxylin-positive nuclei (representing the total amount of nuclei in the image) were counted from the H-channel using the particle-counting algorithm and compared with the thresholded positively immune-stained nuclei counted from the DAB channel. The results of the densitometric image analysis of the serial images for each sample were first averaged, and these single values were then used for the further combined analysis of results. The image analysis macros are available from the authors by request.

The primary outcome measures were safety in terms of hemodynamic adverse effects (ventricular arrhythmias and death) and feasibility of the therapy administration in a clinical setting. Feasibility was evaluated by the success in completing the delivery of the cell sheet to the infarction area in myocardium and in success in closing the right RAA without any additional suturing. Additional outcome measures were changes in LV wall thickness and in LVEF measured by echo, any problems related to recovery such as infection, lack of normal growth, eating or exercise and the inflammatory response to ECM.

The groups were compared, and analyses were performed with SPSS software version 16.0 (Chicago, IL, USA). Two-tailed t-test for independent samples was used to compare the groups. Differences were considered significant at P<0.05. Unless otherwise specified, data in the manuscript are presented as mean**±**SD.

## Results

Ten pigs were operated in total. Two pigs died at the table due to ventricular fibrillation after the LAD ligation. Eight survivors went through the whole study protocol of three weeks. All of them recovered the operation well, without observable changes in normal physical activity or growth. One animal of the control group had a local abscess beneath the sternal wound while reopening the sternum at the three weeks’ time point. Abscess was located below xiphoid process and was not in contact with the heart or the transplant. There was no further evidence of mediastinitis or sternal infection. This animal was physically in good health without any disturbances in appetite or growth. Other animals showed no signs of wound or other infections.

### Feasibility

All eight infarct survivors received the patch successfully. During the observation time of two hours after the ligation, patch was ready to be placed in all cases. There was no bleeding in appendages and no additional patching or suturing was needed. The chosen minimum size for the removable appendage tissue was 5×10 mm, and the weights ranged 610-830 mg.

### Echocardiography

For all pigs, the echo LVEFs evaluated at baseline prior to surgery were normal (LVEF was 70% in all animals). There were no anatomical abnormalities or congenital deficiencies. Each animal’s postoperative echo showed hypokinetic area at three weeks after surgery, but there were no differences in follow-up LVEFs between the groups (AAMs patch group EF 63.3%±9.4%, range 50%–70%; ECM patch group EF 62.5%±4.3%, range 60%–70%, p=0.915).

The mean LV wall thickness (measured at the infarction area) showed no differences between the groups (at baseline AAMs patch group 6.9mm**±**0.6mm, range 5.6mm–7.3mm; ECM patch group 6.7mm**±**0.7 mm, range 5.7mm–7.4mm, *p* = 0.731 and at follow-up AAMs patch group 7.0mm**±**1.8mm, range 4.6mm–8.9mm; ECM patch group 6.8mm**±**0.4mm, range 6.4mm–7.4mm, *p* = 0.881). Thickening of the LV wall during the follow-up was similar between the groups (AAMs patch group 0.3mm±1.7mm, range −2.1mm–1.7mm; ECM patch group 0.1mm±0.6mm, range −0.85mm–0.7mm, *p* = 0.889). One animal from the AAMs patch group had severe bleeding during re-sternotomy and the possibility to provide a comparable echo was lost.

### Histology

HE-staining showed infiltration of inflammatory cells and foreign body reaction, in all animals. This reaction was significantly less in hearts with AAMs, when compared to the ECM hearts (**Fig 2**). The average distance between the Cormatrix^®^ and epicardium was significantly shorter in the AAMs patch group (AAMs patch group 1,463μm**±**441μm, range 659 μm–2,523μm; ECM patch group 2,457μm±865μm, range 1,599μm–4,729μm, *p* = 0.001) and the number of inflammatory cell nuclei was significantly less in the AAMs patch group (AAMs patch group 6682/mm^2^±2,475/mm^2^, range 3,046/mm^2^–11,609/mm^2^; ECM patch group 8,736/mm^2^**±**2,798/mm^2^, range 5,137–13,506/mm^2^, *p* = 0.026) (Figure 2). In the ECM patch group, nuclei were clustered, whereas in the AAMs patch group, more diffuse pattern of the nuclei was observed (**Fig 2**). In both groups, similarly, the inflammation was evident by migration of giant cell macrophages, neutrophils and lymphocytes as characterized by HE-staining. Also, there were plenty of capillary vessels along the foreign body reaction without difference between the groups. The inflammation area in all hearts was limited to the area of cell graft or control graft transplant and the endocardial area showed no inflammation. The fibrotic changes due the infarct were relatively small and mainly seen at the endocardial part of the LV wall. The size of the infarction was variable in both groups and all animals. Only a few samples (from three animals) showed larger infarct with granulation tissue and small calcification and necrotic areas with no difference between the groups.

**Figure 2.**
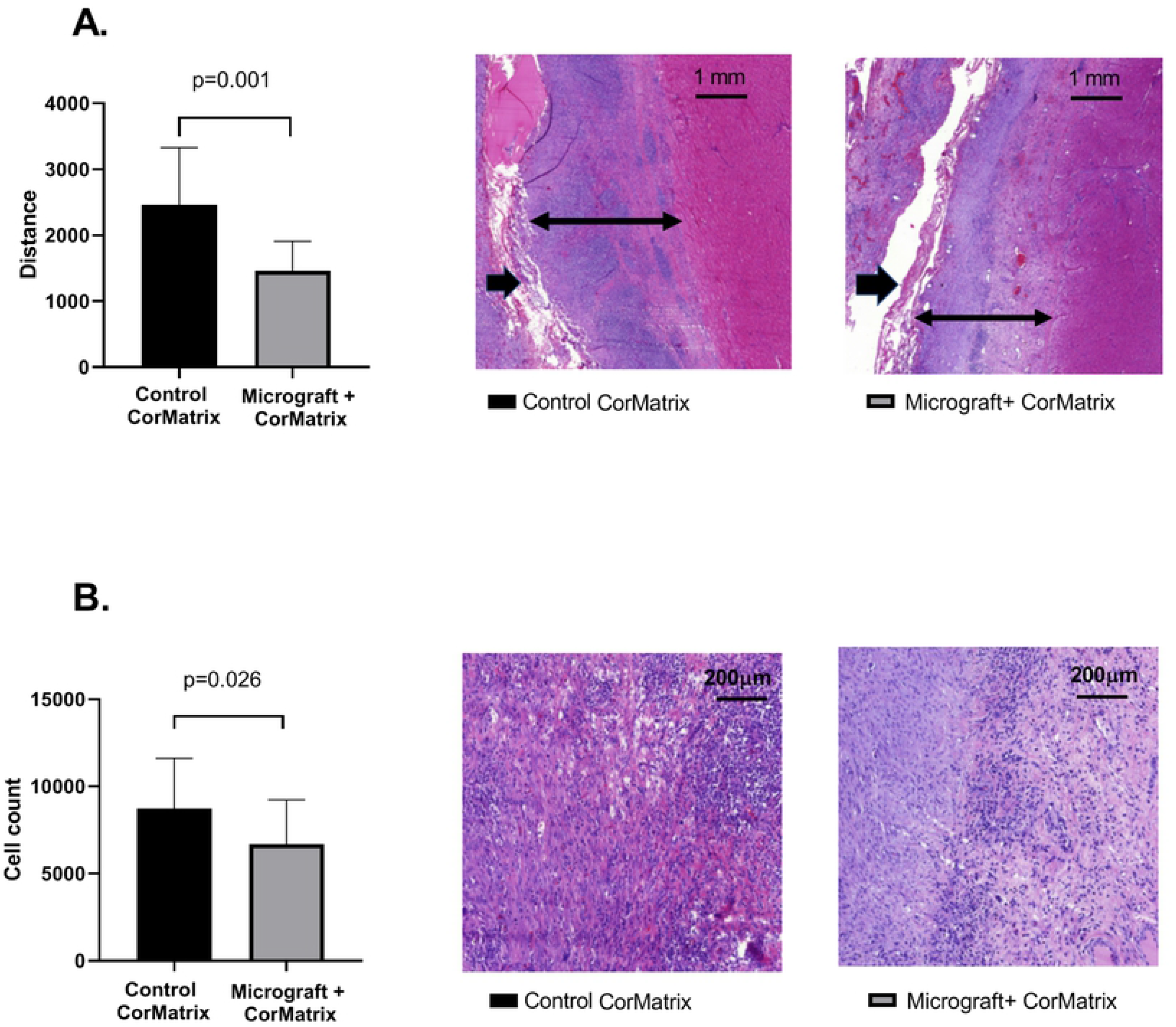
Comparison of inflammatory reaction by hematoxylin eosin staining. Hematoxylin eosin staining from the area of the ECM patch (marked with single arrow) and myocardium. **A.** Figure shows average distance from the ECM patch to epicardium (average of 10 measurements). A representative distance measurement shown with double red arrow. Autologous atrial appendiceal micrograft (AAM) containing samples were significantly thinner than samples without AAMs. Scalebar 1 mm. **B**. Figure showing cell count from the foreign body reaction caused by the patch. Inflammation was more cell dense in control samples whereas the AAM group showed more diffusely scattered nuclei at the supra-epicardial area. The cell nuclei per 1 mm^2^ count was performed twice from each sample. Scalebar 200 μm.

Results from immunohistochemistry analysis of the inflammatory infiltrate are presented in **Table 1**. CD3^+^ cell density, representing the T-lymphocyte density, was greater in AAMs patch group (AAMs patch group 4,834/mm^2^±1,271/mm^2^, range 3,028/mm^2^–7,131/mm^2^; ECM patch group 3,364/mm^2^**±**667/mm^2^, range 2,336/mm^2^–5,346/mm^2^, p < 0.001) (**Fig 3**). Moreover, mean intensity was stronger in the AAM patch group in CD3 staining (AAMs patch group 126/mm^2^**±**12/mm^2^, range 107/mm^2^–146/mm^2^; ECM patch group 114/mm^2^**±**9. 8/mm^2^, range 87/mm^2^–124/mm^2^, p=0.007). The density of CD45^+^ cells, representing total lymphocyte density, was not significantly different between groups (AAMs patch group 6,376/mm^2^**±**1,990/mm^2^, range 4,009/mm^2^–14,748/mm^2^; ECM patch group 5,467/mm^2^**±**1,579/mm^2^, range 3,159/mm^2^–9,925/mm^2^, p = 0.078) (**Fig 4**). Mean intensity in CD45 staining did not reach statistical difference between groups. There were no differences in the CD20^+^ cell densities between groups (**Fig 5**).

**Table 1.** Results and comparison of nuclei from immunohistochemistry stainings. CD3 staining is representative of the T-lymphocyte population, CD45 of the overall lymphocyte population, and CD20 of the B-lymphocyte population.

| CD3 | Cell | Cormatrix |
| --- | --- | --- |
| Positively stained cells/mm<sup>2</sup> | 4,838±1,271 (3,028–7,131) | 3,364±667 (2,336–5,346) |
| Mean staining intensity | 126±12 (107–146) | 114±9.8 (87–124) |
| Stained area/total area (%) | 26±6.0 (15–34) | 21±7.0 (7.9–32) |
| CD45 |  |  |
| Positively stained cells/mm<sup>2</sup> | 6,376±1,990 (4,009–14,748) | 5,467±1,579 (3,159–9,925) |
| Mean staining intensity | 140±21 (109–192) | 129±13 (110–151) |
| Stained area/total area (%) | 42±4.5 (34–50) | 42±7.4 (25–54) |
| CD20 |  |  |
| Positively stained cells/mm<sup>2</sup> | 700±277 (346–1,069) | 682±389 (238–1,552) |
| Mean staining intensity | 137±9.7 (124–152) | 146±29 (91–198) |
| Stained area/total area (%) | 17±5.4 (6.9–24) | 15±7.9 (3.9–33) |

**Figure 3.**
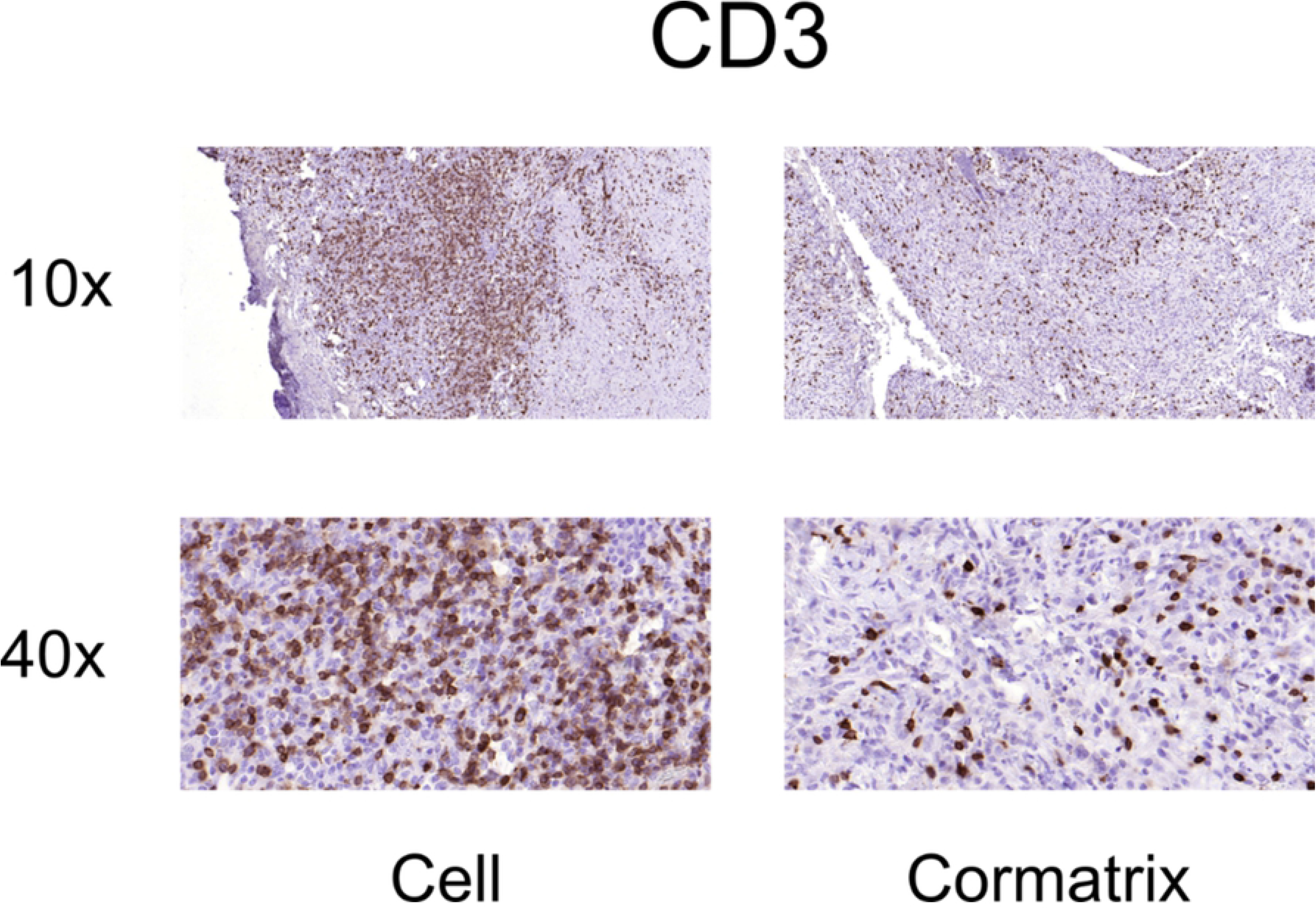
Representative immunohistochemistry images for T-lymphocytes (CD3+) from the AAMs patch group (left panels) and ECM patch group (right panels).

**Figure 4.**
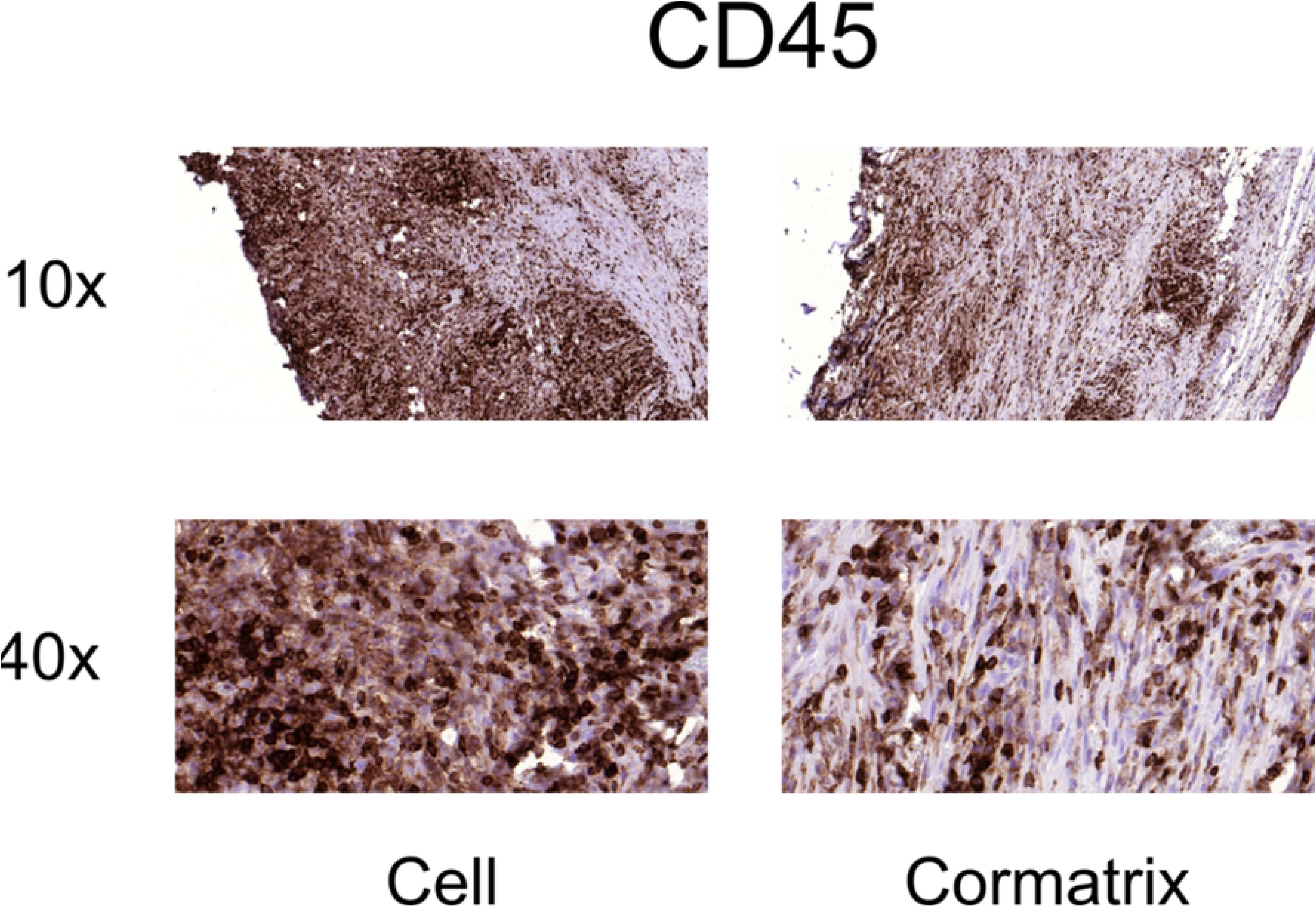
Representative immunohistochemistry images for leukocytes (CD45+) from the AAMs patch group (left panels) and ECM patch group (right panels).

**Figure 5.**
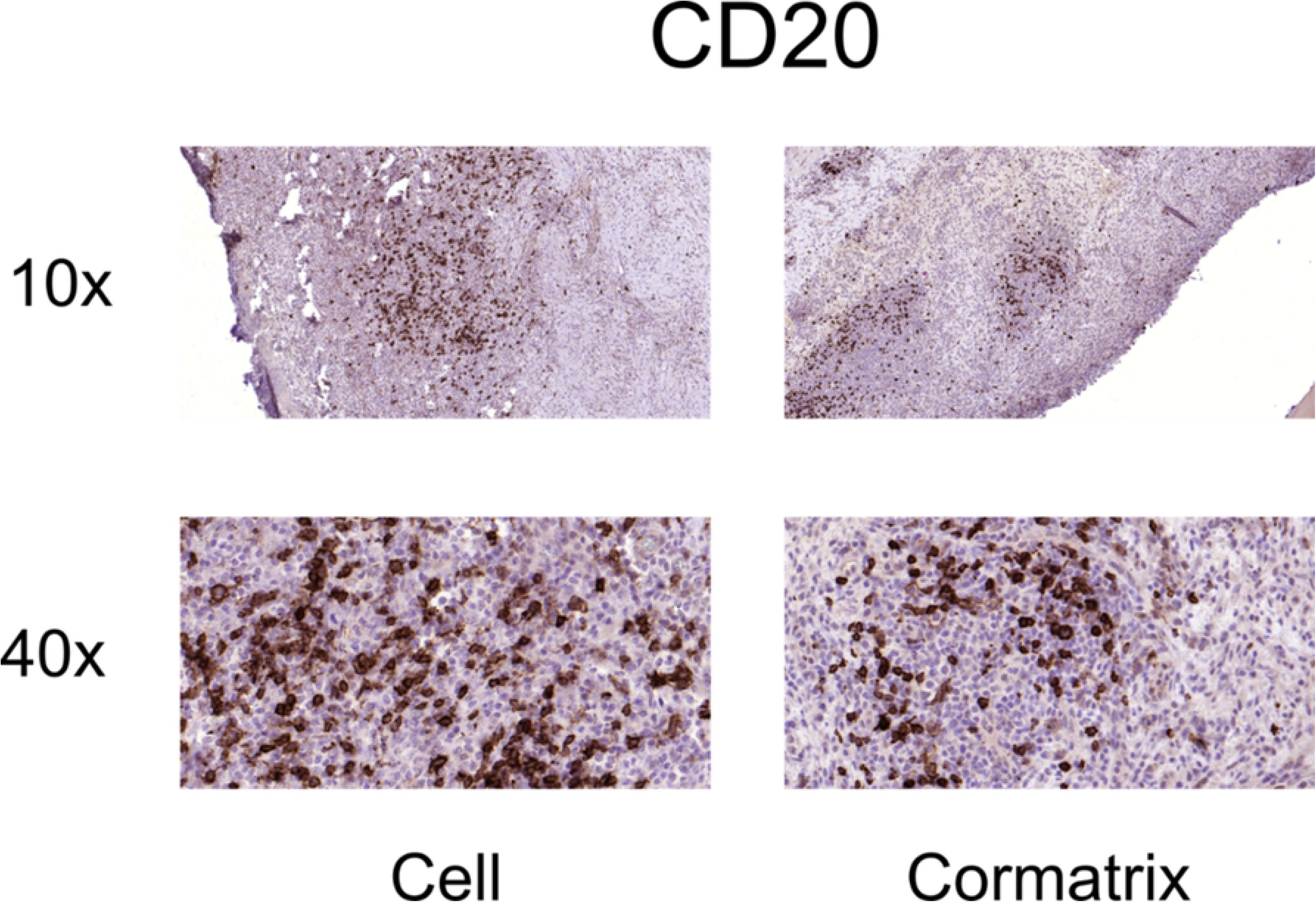
Representative immunohistochemistry images for B-cells (CD20+) from the AAMs patch group (left panels) and ECM patch group (right panels).

## Discussion

The specific aim of this study was to evaluate the safety and feasibility of epicardial delivery of AAMs in pigs after myocardial infarction. We found that epicardial transplantation was both feasible and safe using this large animal model.

Ventricular fibrillation occurred in two out of ten pigs (20%). These arrhythmias occurred immediately after ligation of the LAD and caused death of both animals. The reported mortality rates in porcine models of infarction are usually high, 33% [27] due to the susceptibility of pigs developing fatal arrhythmias after coronary occlusion. The infarction in our study was therefore not too large to cause death but adequate to cause local changes and clearly observable acute ischemia.

All eight pigs receiving therapy recovered well and survived the three-week follow-up. Even though the hypokinetic area in the inferior and anterior wall of LV was seen in echo, the infarction changes seen in histology were not constantly observable in every section. In one histological plate cross section of the left descending artery was seen re-opened for blood flow due to the collateral connections. The natural capacity of these adolescent pigs to recover was most likely the reason for only little permanent damage caused by the infarction. [28]

In all samples, an active foreign body reaction was detected with leukocytes including macrophages, eosinophils and lymphocytes. There was, however, a significant difference in the amount of inflammation among the two groups. The number of inflammatory cell nuclei in the AAMs patch group samples was significantly less as measured by counting the number of nuclei and the area foreign body reaction when compared to the ECM patch group. The cells and micrografts inside the transplant were clearly protecting against the foreign body reaction. This may be explained by the paracrine effect of transplanted cells. Transplanted cells release protective factors and signals that have been proven to be the main reason for environmental tissue repair and remodeling. These signals (including cytokines, chemokines, growth factors, microparticles) activate pathways which consecutively improve blood perfusion, tissue repair and remodeling and inhibit hypertrophy and fibrosis. [29–31] One of the most crucial factors in cell signaling and stem cell derived communication with myocardium is exosome, nanometer-sized vesicle. Exosomes have been proved to increase cell proliferation, reduce infarct size and increase EF in animal studies. [32, 33] In addition these cardio-protective secreted microparticles that induce paracrine and autocrine phenomenon have also been thought to inhibit unwanted inflammation [3–5] via, for example, activation of macrophages, cell apoptosis and increase in metabolism [3, 5]. This protective force against inflammation that transplanted cells possess may be the reason why the inflammation reaction was decreased when micrografts were used compared with the ECM sheet alone.

Cormatrix^®^, decellularized porcine small intestine submucosa, is clinically used as a scaffold to support tissue regrowth in vascular structures. This extracellular matrix is gradually replaced, leaving behind functional tissue. Several studies have shown that Cormatrix^®^ elicits inflammatory reactions [34–36] as also clearly seen in our study. Inflammation is histologically seen mainly in eosinophilic infiltrates often with granulation tissue, fibrosis [36] and foreign body giant cell reaction [37]. Therefore, knowing the inflammatory response that Cormatrix^®^ causes, especially in porcine, the strength of foreign body reaction compared to the use of AAMs and ECM matrix alone was the most important finding and the main focus in histologic evaluation. The use of AAMs seemed to inhibit the inflammatory effect of Cormatrix^®^ and therefore result in a more satisfying environment.

It has been proven that the use of biological material alone may have a beneficial effect to the infarcted myocardium. This is possible due to the fact, that the material placed on myocardium seems to thicken the wall, reduce wall stress and eventually prevent the negative LV remodeling. [38] This was also seen in our study. There were no differences in LV wall measures or LVEF between the two groups. The infarcted wall was performing equally well whether the micrografts were or were not used with the supportive matrix.

Our results demonstrate that atrial micrografts can modulate and suppress the immune response, as mimicked here by the foreign body response to biomaterial, by shifting the CD45+ leukocyte balance towards a more CD3+ T-cell-populated one. These results have important implications for the treatment of acute inflammatory reactions, for example acute myocardial infarctions, with atrial appendage micrografts. Our results provide the first insight into the regulation of the inflammatory and immune responses by atrial appendage micrografts. Further investigations should be targeted towards the identification of leukocyte polarization and especially the anti-inflammatory subtype analysis of leukocytes and T-cells. Moreover, these experiments should utilize various inductors of inflammation and immune responses.

## Conclusions

In conclusion, ECM sheet provides substantial inflammatory response to adjacent tissues, while myocardium stays intact in this response. Additional delivery of AAMs attenuates the inherent inflammatory response, while keeping the regenerative potential as effective, providing a clinically feasible additive for the goal to repair the infarcted myocardium. Transplantation of AAMs with ECM shows good clinical applicability as adjuvant therapy to cardiac surgery and may serve as a potential delivery platform for future cell or gene-based cardiac therapies.

## Acknowledgements

Research nurses Liisa Blubaum, Anna Blubaum and Mariitta Salmi are gratefully acknowledged and thanked for their professional expertise in the development of the protocol. Technicians Olli Valtanen and Veikko Huusko are gratefully acknowledged and thanked for their help and professional contribution in animal laboratory.

